# Distribution and asymptotic behavior of the phylogenetic transfer distance

**DOI:** 10.1101/368993

**Authors:** Miraine Dávila Felipe, Jean-Baka Domelevo Entfellner, Frédéric Lemoine, Jakub Truszkowski, Olivier Gascuel

**Affiliations:** Unité Bioinformatique Evolutive / C3BI USR 3756, Institut Pasteur & CNRS, Paris, France; Biosciences eastern and central Africa (BecA-ILRI Hub), International Livestock Research Institute, PO Box 30709, Nairobi 00100, Kenya; Hub Bioinformatique et Biostatistique, C3BI USR 3756, Institut Pasteur & CNRS, Paris, France; Méthodes et Algorithmes pour la Bioinformatique, IBC - LIRMM UMR 5506, Université de Montpellier & CNRS, Montpellier, France

## Abstract

The *transfer distance* (TD) was introduced in the classification framework and studied in the context of phylogenetic tree matching. Recently, Lemoine et al. (2018) showed that TD can be a powerful tool to assess the branch support of phylogenies with large data sets, thus providing a relevant alternative to Felsenstein’s bootstrap. This distance allows a *reference branch β* in a reference tree 𝒯 to be compared to a branch *b* from another tree *T*, both on the same set of *n* taxa. The TD between these branches is the number of taxa that must be transferred from one side of *b* to the other in order to obtain *β*. By taking the minimum TD from *β* to all branches in *T* we define the *transfer index*, denoted by *ϕ*(*β, T*), measuring the degree of agreement of *β* with *T*. Let us consider a reference branch *β* having *p* tips on its light side and define the *transfer support* (TS) as 1 – *ϕ*(*β, T*)*/*(*p –* 1). The aim of this article is to provide evidence that *p* − 1 is a meaningful normalization constant in the definition of TS, and measure the statistical significance of TS, assuming that *β* is compared to a tree T drawn according to a null model. We obtain several results that shed light on these questions in a number of settings. In particular, we study the asymptotic behavior of TS when *n* tends to ∞, and fully characterize the distribution of *ϕ* when *T* is a caterpillar tree.

## 1 Introduction

The *transfer distance* or *R-distance* was introduced in the classification framework by Day [5] and Régnier [22], as a measure of (dis)similarity between partitions of a set. It is defined as the minimum number of elements that need to be removed from their original class or transferred from one class to another, in order to transform one partition into the other. This distance possesses some desirable properties, for example its low computational cost in comparison with other metrics, as established by Day [5]. Charon, Denœud, Guénoche, and Hudry [4] studied other characteristics of this distance such as the maximum transfer distance that can be obtained when comparing two partitions with a fixed, but possibly different, number of classes. As highlighted by Denœud [7], it proves challenging to study the theoretical properties of the transfer distance, so the author proposed an experimental analysis of the transfer distance using simulations, to discuss its interpretation and approximate its distribution and mean when considering pairs of random partitions of the same set.

The interest in using the transfer distance to compare phylogenetic trees started with a seminal paper by Day in 1985 [6]. In the field of computational biology, problems involving tree comparison have remained a major challenge for many years. A common concern with most of these problems is to define a suitable metric on trees. The transfer distance is a measure to compare bipartitions, and a phylogenetic tree is unambiguously defined by the set of bipartitions induced by its branches. Then, a logical question to ask is whether we can define a metric on trees based on this transfer distance on bipartitions. There is not a unique way of defining such a metric and several authors have worked on the subject. For instance, in [6], the author proposes several algorithms and methods to solve related tree problems, in particular the construction of the consensus of a set of trees. As discussed in [6], this task requires the optimization of a consensus index, which can be defined using the transfer distance or other metrics, such as the well-known Robinson-Foulds (RF) metric [21]. The latter is probably the most widely used distance between trees and is defined as the number of bipartitions belonging to one tree but not to the other. However, the RF metric is known to have several drawbacks, including its lack of robustness, since it is highly sensitive to certain small tree changes, as pointed out by Lin, Rajan, and Moret [19] and Bogdanowicz and Giaro [3].

In another study, Boc, Philippe, and Makarenkov [2] proposed to optimize a tree comparison index that can also be based on metrics, including RF and the transfer distance. Having set a goal to detect accurately horizontal gene transfer events, the authors showed that the version of their algorithm relying on the transfer distance provides the best results when searching for an optimal scenario of Subtree Pruning and Regrafting (SPR) moves needed to transform a gene tree into a species tree. However, the transfer-based dissimilarity defined in [2] is not a metric, since it violates the triangle inequality in some cases [2, p.197, Prop. 1]. Lin, Rajan, and Moret [19] addressed this problem by proposing a different distance measure based on minimum-cost matching between the two sets of splits induced by both trees. For that metric, also relying on the transfer distance, the triangle inequality holds. Additionally, the computational and statistical properties of this new distance are studied in [19], where the authors provide a low-polynomial time algorithm, establishing its robustness through statistical testing and demonstrating its usefulness in clustering trees.

Recently, we proposed a new bootstrap method for large phylogenetic trees, relying on branch comparisons based on the transfer distance [18]. The aim of that study was to use a more fine-grained measure for the presence of a branch in a tree, rather than the binary values used in Felsenstein’s classical bootstrap technique [11]. We compared a *reference branch β* in a *reference tree* 𝒯 to another tree *T*, typically a bootstrap tree, by taking the minimum of the transfer distance from *β* to any branch *b* in *T*, which we called the *transfer index* and denoted by *ϕ*(*β, T*). Next, we averaged the values of *ϕ* over a set of bootstrap trees, obtaining, after appropriate normalization, the so-called *transfer bootstrap expectation* (TBE). We explored the behavior of TBE as a measure of support for the branches of a phylogenetic tree, compared to that of Felsenstein’s support (FS). In a number of experiments using both real and simulated data, we found that TBE outperformed FS. This was particularly noticeable for deep branches and large values of *n*, where FS often failed to detect the phylogenetic signal in the trees. In view of those results, TBE shows promise as a useful tool in phylogenetic analysis. In [18], we studied and discussed several of its mathematical properties, but there is still need for further work so that the transfer index and TBE are fully understood. The main motivation for the present work is therefore to study the properties of the transfer index and support, assess their statistical significance, and give analytic expressions for TBE when the reference branch is compared to a tree *T* drawn randomly according to some null model.

To be more specific about the results obtained here, fix *n ≥* 4 and let us consider phylogenetic trees on a set *X* of *n* taxa. To distinguish the two sides of the reference bipartition *β*, we say that its *light side* contains *p ≥* 2 taxa while its *heavy side* has *n − p ≥ p* taxa. The TBE we proposed in [18] is actually the average, over all the bootstrap trees, of the *transfer support* function (TS), which is defined as 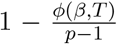. It is not hard to see that *ϕ*(*β, T*) *≤ p −* 1 and thus the TS function takes values on the interval [0, 1]. Notice that we deliberately exclude here cases with *n* = 2, 3, and *p* = 1, which result in trivial bipartitions.

First, let us reproduce the results obtained from computer simulations in [18], where we looked at four different models for the tree *T*. More precisely, we considered (1) two simple models for the topology of phylogenetic trees: caterpillar trees and perfectly balanced binary trees; and (2) two classic null models for speciation: the Proportional to Distinguishable Arrangements (PDA) and Yule-Harding models (see Section 2 for further details). For each of these models, we performed simulations for different values of *n* (128, 256, 512, and 1,024) and all the possible values of *p* for each *n* (i.e. 2 *≤ p ≤* ⌊*n/*2⌋). Then, for each value of (*n, p*) and for each model, we simulated a set of reference bipartitions and a set of trees to be compared to these reference bipartitions. The results are displayed in Fig. 1, where we plotted the TS values thus obtained against the different values of *p*. This figure shows striking evidence that, on average, TS stays close to 0 with random trees, so the transfer index stays close to its upper bound *p −* 1 for all *p*. Additionally, we observe that the maximum value of the TS attained over all possible values of *p* seems to be obtained at *p* = ⌊*n/*2⌋ and to decrease when *n* increases. Moreover, this asymptotic behavior seems to be independent of the topology/model considered for the tree *T*. These initial observations motivate the questions we address here.

**Figure 1:**
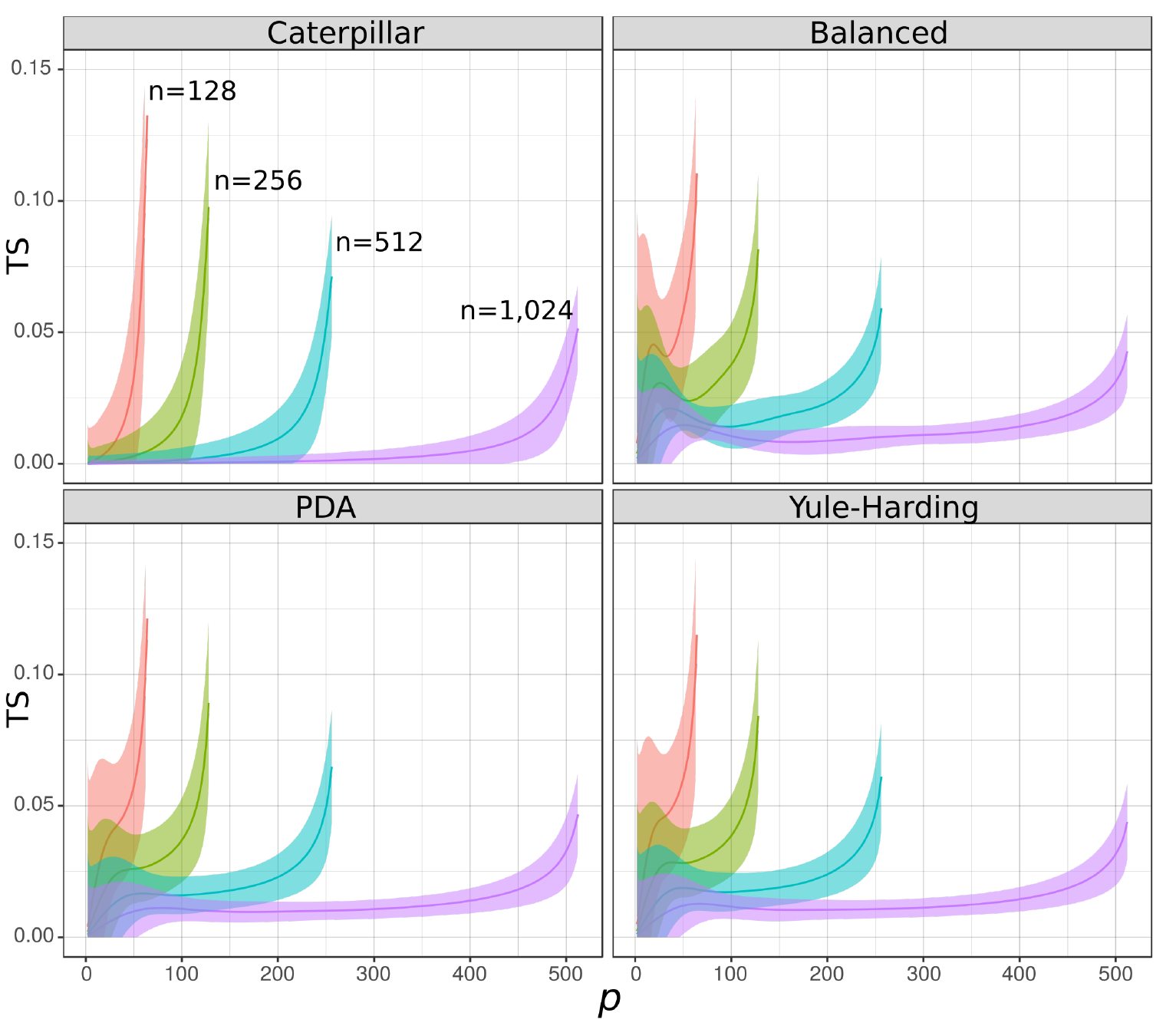
Simulations of TS for random trees. The four panels correspond to the four different models considered: caterpillar, totally balanced, PDA, and Yule-Harding. For each of these models, we considered four different values of *n* and used the following color codes to distinguish them: 128 (red), 256 (green), 512 (blue), and 1,024 (purple). For each model and for each value of *n* and *p* (from 2 to ⌊*n/*2⌋), we simulated 100 reference bipartitions and 1,000 trees to be compared to the reference bipartition. We plotted the mean of the 100 *×* 1, 000 TS values obtained for each *n* and for each model (bold lines) and the standard deviation (shaded areas), against *p*.

The intuition provided by computer simulations led us to obtain several theoretical results that justify what we observed in Fig. 1 and the simple normalization by *p −* 1 used in [18]. We also formulate a set of conjectures that are discussed in Section 6. Our first result consists in the characterization of the asymptotic behavior of the transfer index *ϕ*(*β, T*) for a random bipartition *β* of the set *X* and any tree *T*. More precisely, we prove that the transfer index converges in probability to *p −* 1 when *p* is fixed (but see below) and *n* tends to *∞*. The proof relies on the comparison of the transfer index with the parsimony score of a binary character and the use of a result from Steel and Penny [25]. We then use concentration inequalities to characterize the asymptotic behavior of the transfer index when *p* grows, depending on *n* (e.g. when *p →* ⌊*n/*2⌋). Lastly, when *T* is a caterpillar tree, we fully characterize the probability distribution of the transfer index based on a one-to-one correspondence between these trees and North-East (NE) lattice paths, a common technique for counting combinatorial objects [20]. All of these results show that *p −* 1 is the appropriate normalization constant for the TS proposed in [18] and that this support has the expected behavior, which is that, in the absence of phylogenetic signal in the tree *T* regarding the reference branch *β*, the TS is close to 0, especially for large trees.

The paper is organized as follows. In Section 2, we give the main definitions and properties of the concepts described earlier. Section 3 is devoted to the results concerning the parsimony score, and Section 4 presents the asymptotic results using concentration inequalities. Details on the specific case of the caterpillar tree are given in Section 5.

## 2 Preliminaries

In this section, we give the main definitions and general properties on phylogenetic trees that are needed for the rest of the paper. We refer to [23] for an extensive mathematical treatment of this subject.

Let us fix *n ≥* 4 and *X*, a set of *n* taxa. We consider phylogenetic trees on *X*, that is trees whose leaves are mapped one-to-one to *X*. These trees are called *phylogenetic X-trees* or simply phylogenies. For simplicity of notation, we shall always take *X* = {1, 2*, …, n*}. Denote by UB(*n*) the set of all unrooted *binary* phylogenetic trees (every interior vertex has degree 3) on *n* leaves. For a phylogenetic tree *T*, we use 𝜀(*T*), *𝓔*(*T*) to denote respectively the set of edges and the set of vertices of the tree.

For any *X*-tree *T*, a branch *b* ∈ 𝜀(*T*) can be encoded in several equivalent ways, that we will use indistinctly depending on the context. First, any branch *b* defines a bipartition (or split), and we can associate *b* to a vector *υ*(*b*) in {0, 1}^*n*^ by assigning the same number (e.g. 0) to all the elements on the same side of the split induced by this branch. Notice that *b* is also encoded by *υ*̅(*b*), the negation of *υ* (i.e. the 0 values are turned into 1 and vice versa). Likewise, we can identify a bipartition, with a *bicoloration* of the leaves, that is a function that assigns one of two colors (black = **B** or white = **W**) to each leaf label. Notice however, that we can consider a bipartition or a bicoloration on a tree that does not correspond to any branch in this tree. To make the distinction, we say *b* ∈ 𝓔(*T*) for the bipartitions induced by branches on the tree *T*, and we use 𝓧 : = {*f* : *X →* {**B**, **W**}} to denote the set of all possible bicolorations of the tips of *T*. Then, *b* ∈ 𝓧 does not necessarily correspond to a branch in *T*, but to a bicoloration of its tips.

We will leverage the visual aspect of the two-color representation and, throughout the rest of the article, we associate the *p* taxa of the light side in the reference bipartition with the black color **B** and the *n − p* (*≥ p*) taxa of the heavy side with the white color **W** (Fig. 2, left). The set of bicolorations satisfying that *|*{*i* ∈ *X* : *f* (*i*) = **B**}*|* = *p* will be denoted by 𝓧_*p*_. A tree *T* endowed with a bicoloration is called a bicolored tree.

**Figure 2:**
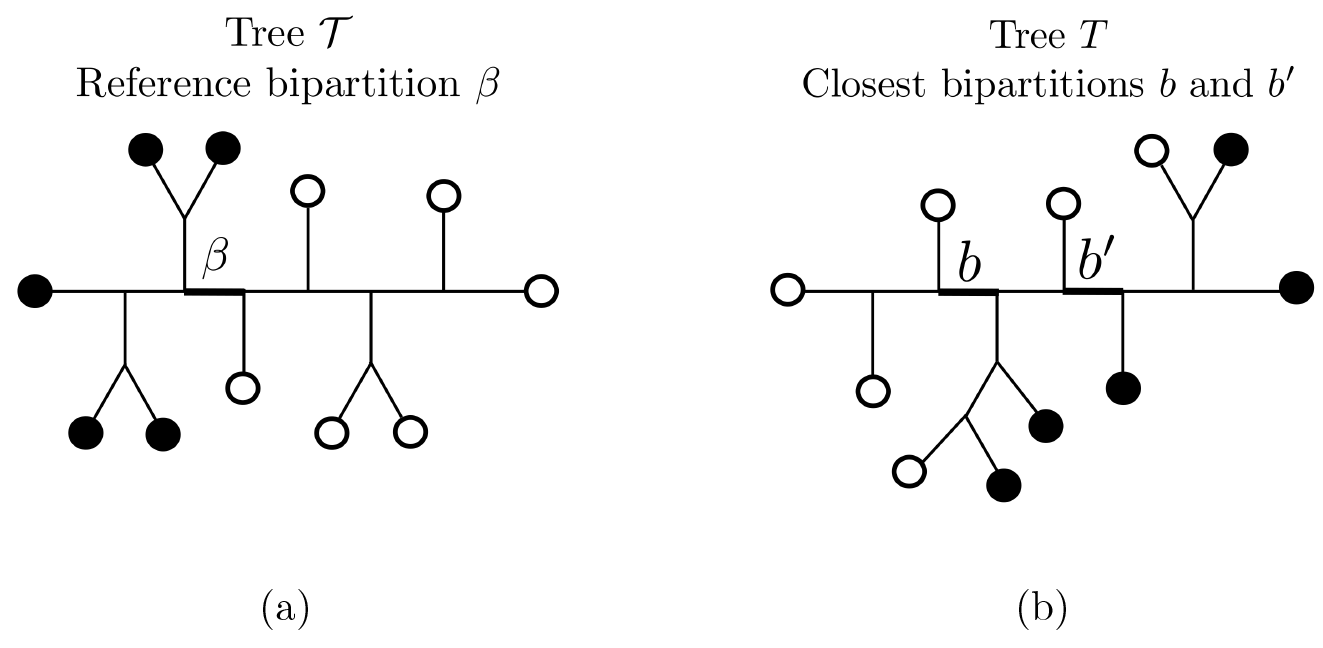
(a) An example of reference branch *β* dividing *X* in *p* = 5 black tips and *n – p* = 6 white tips, respectively. (b) Another *X*-tree *T* in which the closest bipartitions to *β* are *b* and *b′*, both giving a transfer distance *δ*(*β, b*) = *δ*(*β, b′*) = 3, and thus *ϕ*(*β, T*) = 3 as no other branch in *T* is closer to *β*.

As described in the Introduction, the transfer distance is used to compare a branch *β* in the reference phylogenetic tree topology 𝒯, to a second branch *b* in another tree topology *T*, both on the same taxa set *X*. This distance can easily be defined using the Hamming distance *H*(*·, ·*) between two vectors of equal size.

### Definition 1 (Transfer distance)

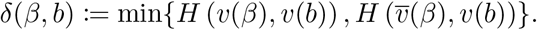

Based on this definition, notice that *δ*(*β, b*) = 0 if and only if *υ*(*β*) and *υ*(*b*) define the same bipartition. To measure the degree of presence of *β* in *T*, we define the *transfer index*, denoted by *ϕ*(*β, T*), which is the minimum of the transfer distance over all branches in *T* [18].

### Definition 2 (Transfer index)

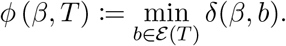

As mentioned before, we are interested in the case where the reference tree 𝒯 and a branch *β* on this tree are fixed. The core idea in [18] is to measure the presence of this reference branch in a set of bootstrap trees by using the following *transfer support* function.

### Definition 3 (Transfer support)

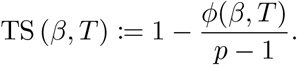

The transfer index and support functions satisfy simple properties that we can find in [18], and that are included here for completeness:

(i) *ϕ* (*β, T*) = 0 ⟺ *β* ∈ *T*,
(ii) *ϕ* (*β, T*) ∈ [0*, p −* 1] or equivalently TS (*β, T*) ∈ [0, 1].

The first statement can be deduced directly from the definition of *δ*(*β, b*), which is 0 if and only if *υ*(*β*) = *υ*(*b*) or *υ*̅(*β*) = *υ(b*). Thus, *ϕ* (*β, T*) = 0 if and only if we can find the bipartition induced by *β* in *T*. Moreover, we say that a bipartition is trivial when it has a single leaf on one side, and the remaining *n −* 1 leaves on the other. For a trivial bipartition *b* defined by a taxon that belongs to the light side in *β*, we obtain that *δ*(*β, b*) = *p −* 1, and since *ϕ* is defined as the minimum taken over all possible branches in *T*, we obtain the statement (ii).

### Null models

The aim of this study is to characterize the distribution and the asymptotic behavior of the transfer index and transfer support when the reference bipartition *β* is compared to a binary phylogenetic *X*-tree *T* that follows a certain null model. We are interested mainly in unrooted trees, but it should be noted that the existence of a root has no influence on transfer distance values: both branches adjacent to the root define the same bipartition.

There are two ways to define the probabilistic models we are considering. First, we can suppose that we have a fixed bicoloration *χ_p_* ∈ 𝓧_*p*_ and that we draw a tree *T* randomly from UB(*n*), following some specific (probabilistic) model. Another way is to consider that the tree is fixed and a bicoloration of its tips is uniformly chosen from 𝓧_*p*_. In the first case, an interesting question is to consider the probabilistic models that are most commonly used in the field of phylogenetics [14], such as the Yule-Harding or PDA models. On the other hand, for a fixed tree, a natural question is to look at the two extreme cases for the topology regarding balance. The most imbalanced tree is called the caterpillar tree and can be defined as a binary phylogenetic tree for which the induced subtree on the interior vertices forms a path graph (if the tree is rooted, then the root is at one end of the path). On the other side, we have perfectly balanced trees, that is rooted binary phylogenetic trees with *n* = 2^*h*^ leaves (for some *h* ∈ ℕ), each of which is at a distance of exactly *h* edges from the root. We refer to [23, 24] for further details on these tree models.

As explained in the Introduction, we performed computer simulations for these four models to exhibit their asymptotic properties (Fig. 1). We observe that the asymptotic behavior of the TS seems to be independent of the model considered, which we explain in the following sections. Then, a full theoretical treatment is carried out for the caterpillar model.

## 3 Comparing the transfer index to the parsimony score

We are now interested in comparing the transfer index to the widely used parsimony score introduced by Farris [9], Fitch [12], and Hartigan [15]. We then use the result to obtain a first characterization of the asymptotic behavior of the transfer index.

### Definition 4 (Parsimony score)

Consider a phylogenetic tree *T* and a bicoloration of its tips *χ* ∈ 𝓧. Consider an extension of this bicoloration to all the nodes in *T* and denote it by *χ*̅ (each internal node is also assigned one of the two colors). Define

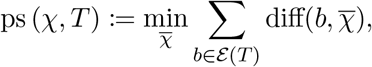

where diff(*b, χ*̅) is 0 if both nodes connected by *b* have the same color in the extension *χ*̅, and 1 otherwise.

By using a simple argument, one can prove the following result from [18], given here for the sake of completeness.

### Lemma 5.

*For any given X-tree T and any bicoloration χ* ∈ 𝓧*, we have that*

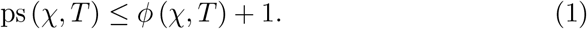

*Proof.* Consider a branch *b* in *T*, and suppose it defines a bipartition having respectively *B_l_*(*b*), *W_l_*(*b*), *B_h_*(*b*), *W_h_*(*b*) black and white tips on its light and heavy sides. We know from the definition that

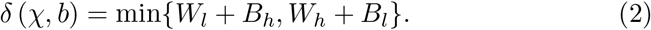

Suppose, without loss of generality, that the minimum is *W_l_* + *B_h_*. We will now look at the parsimony score for this bicoloration. We put a character change at each of the *W_l_* white leaves on the light side of *b* and at each of the *B_h_* black leaves on its heavy side (*parents* take the opposite color). Then the internal nodes on the light side are colored in black and those on the heavy side in white. The number of color changes of this extension is *W_l_* + *B_h_* + 1 because we have to add an extra change at branch *b*. Since the parsimony score of the bicoloration is the minimum taken over all the possible extensions, we have that

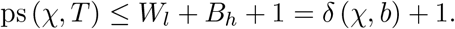

Since this is true for any branch *b* ∈ *T*, we obtain the announced result.

### 3.1 Asymptotic results for fixed *p*

In this subsection, we use inequality (1) between the parsimony score and the transfer index to establish that the transfer index converges to *p−*1 when *p* is fixed and *n* grows to infinity. Let us consider a random bicoloration *χ_p_* from 𝓧_*p*_. Let *T* be any binary tree topology with *n* tips colored by *χ_p_*. The larger *n*, the more dispersed the black tips in *T*, and the higher the probability that the parsimony score is equal to *p* and the transfer index to *p −* 1. This is formalized as follows.

#### Proposition 6.

*Let T be any binary tree topology with n tips, and χ_p_ be a random bicoloration of the tips of T, uniformly chosen from* 𝓧*_p_. We have*

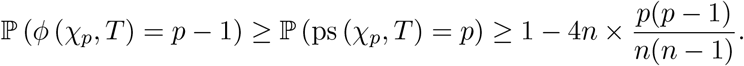

*Proof.* The first inequality is an obvious consequence of inequality (1) and upper bound *p −* 1 on the transfer index. To demonstrate the second inequality, we use a result from Steel and Penny [25, Prop. 9.4.1], stating that if the color (state) changes in a tree are rare enough that any two edges with changes are separated by at least three edges with no changes, then the parsimony score is guaranteed to coincide exactly with the number of changes within the tree. Since *χ_p_* contains *p* black tips, if we color all internal nodes white, we will have *p* changes on the external edges leading to the black tips. If the number of edges separating any pair of two black tips is at least 5, then the parsimony score ps (*χ_p_, T*) is equal to *p*, as all changes are separated by at least 3 internal edges with white vertices at both ends. Thus, we have:

ℙ (ps (*χ_p_, T*) = *p*) *≥* ℙ (any pair of black tips is at distance *≥* 5)

*≥* 1 *−* ℙ (at least one pair of black tips is at distance *≤* 4).

For any tip in the (binary) tree *T*, the number of tips that are at most 4-edge distant is at most 8, and thus the number of pairs of tips at a distance of 4 edges or less is at most 4*n*. The probability of drawing such a pair of tips (among *n*(*n −* 1)*/*2) is

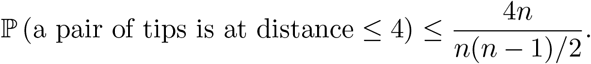

As we have *p*(*p −* 1)*/*2 pairs of black tips, we obtain the desired inequality by the union bound.

This result is valid for any topology *T* as long as the bicoloration *χ_p_* is uniformly distributed in the set 𝓧_*p*_. It has the following immediate consequences.

#### Corollary 7.

*For any tree T, and any χ_p_ uniformly distributed in the set* 𝓧*_p_, we have that,*

- *when p is fixed, the transfer index ϕ*(*χ_p_, T*) *converges in probability to p −* 1 *when n → ∞;*
- *when 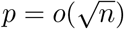, we have that ϕ*(*χ_p_, T*)*−*(*p−*1) *converges in probability to* 0 *when n → ∞.*

## 4 Behavior of the transfer distance when *p* grows with *n*

In the previous section, we showed that, when *n* tends to infinity, the transfer index converges in probability to *p−*1 for fixed *p*, and TS converges to 0 when *p* grows slowly as 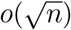. However, simulations in Fig. 1 suggest that the transfer index also behaves in a similar manner for larger values of *p* relative to *n*. For example, for all null models when *p* = ⌊*n/*2⌋, the expected value of TS is larger than 0.1 with *n* = 128, but lower than 0.05 with *n* = 1, 024. In this section, we will show that for “all values of *p*”, the distribution of the transfer index is *concentrated* around *p −* 1, meaning that the probability of the transfer index being “far away” from *p −* 1 vanishes as *n* grows. This explains what we observe in our simulations and motivates the use of *p −* 1 as the normalization term in the definition of TS.

The results we obtain in this section are based on concentration inequalities. More precisely, we make use of the well-known Chernoff-Hoeffding bounds for sums of independent random variables, as stated by Dubhashi and Panconesi [8]. In his original paper, Hoeffding [17] proved that these inequalities also hold for sums of variables obtained by sampling without replacement, which is the case of interest here. The following lemma is a direct consequence of the results in [17] and [8].

### Lemma (Chernoff-Hoeffding bound)

*Let 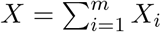 where X_i_ are drawn without replacement from a multiset containing elements between* 0 *and* 1*. Then, for any r >* 0

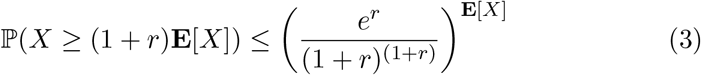

*and for any t >* 0*, we have*

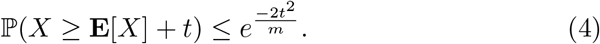

We can now state the main theorem of this section.

### Theorem 8.

*Let T be any binary tree topology with n tips, and let χ_p_ be a bicoloration of the tips in T chosen uniformly at random from* 𝓧*_p_. Then, there exists N* ∈ ℕ*, s.t. for all n ≥ N, with probability at least* 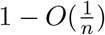,

1. *if p* = *O*(*n^α^*) *for some* 0 *< α <* 1*, then ϕ*(*χ_p_, T*) *≥ p − C for some constant C;*
2. *if p* = *cn* + *o*(*n*) *for some* 0 *< c <* 1*/*2*, then ϕ*(*χ_p_, T*) *≥ p − C* log *n for some constant C;*
3. *if 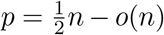, then 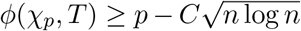 for some constant C.*

These three cases correspond to different growth rates of *p*, from the slowest (1) to the fastest (3). Case 1 is already partly covered for 0 *< α <* 1*/*2 by Proposition 6 and Remark 7, which imply that for these values of *p*,

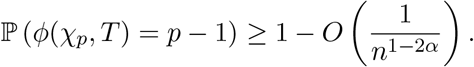

Case 2 corresponds to what we observe in Fig. 1 (see also Extended Data Fig. 1 in [18]), where for any given ratio *p/n* (e.g. *p/n* = 1*/*4), the expected value of the TS decreases when *n* increases. In Case 3, *p* is as large as possible, and the difference between *ϕ* and *p −* 1 is the largest among all three cases, as expected. Note that the bound in Case 3 also holds when *p* = *n/*2 when *n* is even and *p* = *n/*2 *−* 1 when *n* is odd.

### Corollary 9.

*For any tree T, any χ_p_ uniformly distributed in the set 𝓧_p_, and any p that grows with n as in cases 1,2, and 3, the transfer support TS*(*χ_p_, T*) *converges in probability to* 0 *when n → ∞.*

*Proof of Theorem 8.* Consider a bipartition *b* ∈ *T* and let *s* ∈ {*l, h*} denote the light/heavy side of *b*. Let *q_s_*(*b*) be the number of taxa on side *s* of *b* and *B_s_*(*b*) be the random variable corresponding to the number of black taxa on side *s* of *b*. The transfer distance between *b* and the bicoloration *χ_p_* can be written as

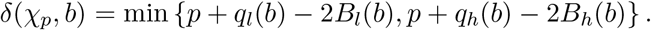

Consequently, we can write the transfer index as

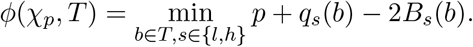

For any 1 *< u < p*, define 𝓑*_u_* = {*b* ∈ *T* : *q_l_*(*b*) *≥ u*}, the set of bipartitions in *T* with at least *u* tips on both sides. We are interested in the set 𝓑_*u*_ since only the bipartitions in this set can give *δ*(*χ_p_, b*) *≤ p − u*. This statement derives from some simple arguments based on the definition of the transfer distance. Consider any *b*′∉ 𝓑_*u*_, we necessarily have *q_l_*(*b′*) *< u < p < q_b_′h*, 0 *≤ B_l_*(*b*′) *≤ q_l_*(*b′*), and 0 *≤ B_h_*(*b′*) *≤ p*, which implies that

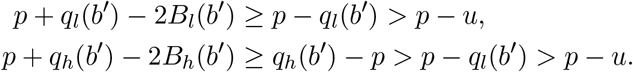

As a consequence, we can bound the tail probability of the transfer index using the union bound over all bipartitions in 𝓑_*u*_, that is

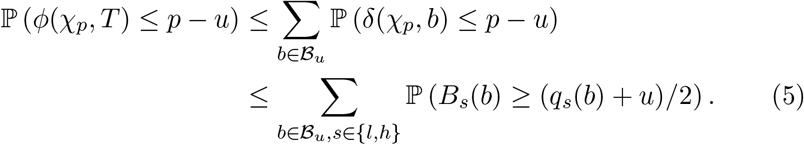

We will now derive a bound on each of the elements of the above sum. First, notice that every *B_s_*(*b*) can be written as

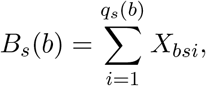

where *X_bsi_* = 1 if the *i*-th leaf on side *s* of *b* is colored black and 0 otherwise. Thus, *B_s_*(*b*) follows a hypergeometric distribution and we have

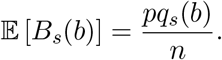

Moreover, Chernoff-Hoeffding inequalities (3) and (4) apply to these hypergeometric variables and enable us to derive the appropriate bounds.

We now must consider three cases depending on the growth rate of *p* with respect to *n*.

*Case* 1*. p* = *O*(*n^α^*) for some *α <* 1.

Applying the bound in (3) with *r* = *n* (*q_s_*(*b*) + *u*) */* (2*pq_s_*(*b*)) *−* 1, we get

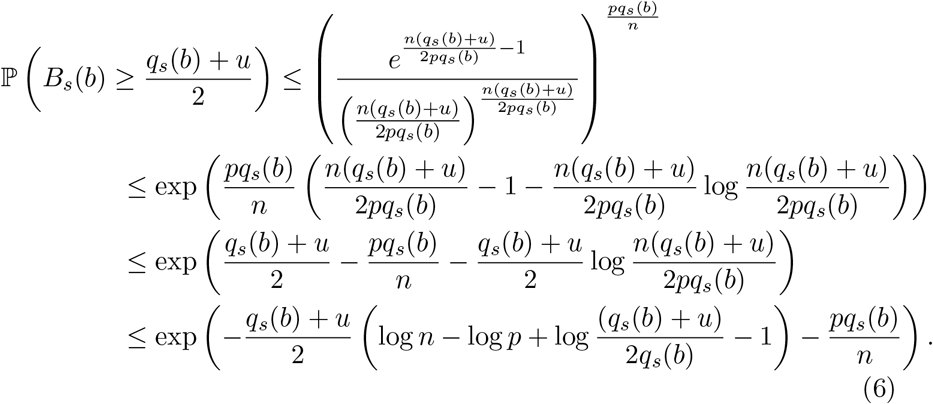

Since *p* = *O*(*n^α^*) for some 0 *< α <* 1, there exist some *N*_1_ ∈ ℕ and *A >* 0 such that *p ≤ An^α^* ∀*n ≥ N*_1_. This implies that for *n ≥ N*_1_, we have log *n −* log *p ≥* (1 *− α*) log *n −* log *A*, which, used in (6), gives

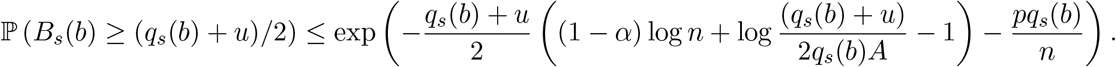

Using the fact that *q_s_*(*b*) *≥ u* for any *b* ∈ 𝓑_*u*_, we get

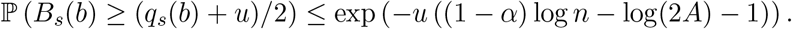

Taking *u* = 2*/*(1 *− α*) and using (5), we get

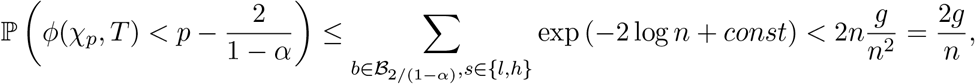

where *g* is a constant and we used the fact that *|*𝓑_*u*_ × {*l, h*}*| <* 2*n* for any *u*. It follows that 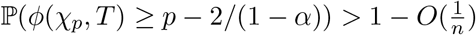 as required.

*Case* 2*. p* = *cn* + *o*(*n*) for some 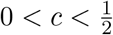.

Applying (4) with 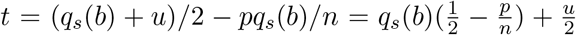, we get

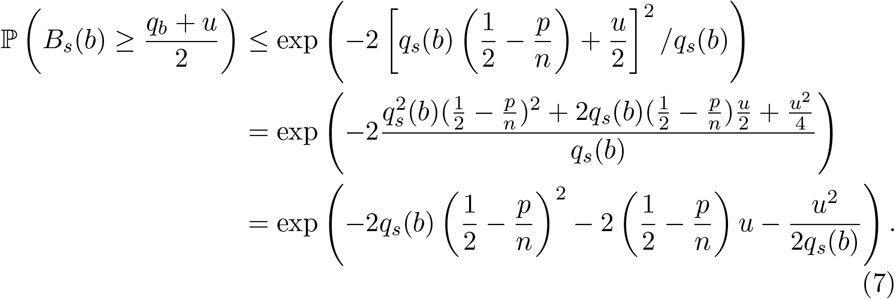

Dropping the last term in the exponent and again using the fact that *q_s_*(*b*) *≥ u*, we get

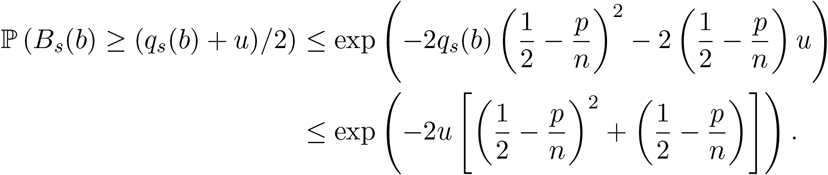

Recall that *p* = *cn* + *o*(*n*). Let *c < c* + *∈ <* 1*/*2. For some *N*_2_ ∈ ℕ, we will have that *p/n < c* + *∈* for all *n ≥ N*_2_, so we can take *u* = *C* log *n*, with 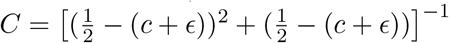 and use (5) to get that

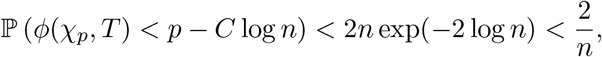

which gives 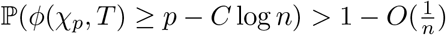 as required.

*Case* 3. 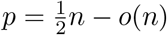

Since 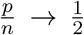 as *n → ∞*, the first two terms in the exponent on the right-hand side of (7) tend to 0, so the bound from the previous case is no longer useful. Based on (7), we can write

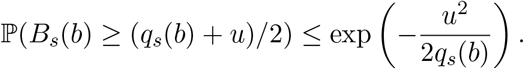

Knowing that *q_s_*(*b*) *< n* for any choice of *b*, we can take 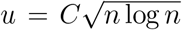, which gives

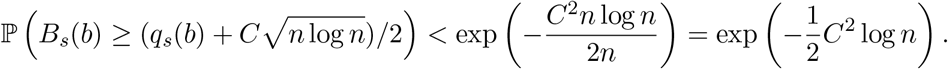

Setting *C* = 2 and using (5), we get

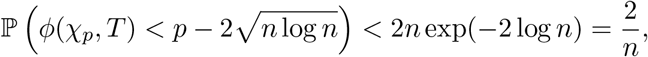

which gives us the result.

## 5 Exact distribution of the transfer index on caterpillar trees

In this section, we provide exact formulae for the transfer index distribution on caterpillar trees. We shall see in the discussion section that these formulae can be used to compute *p*-values for the general case, under suitable assumptions (conjectures). Moreover, the combinatorial techniques used here could potentially help obtain similar results with other trees (e.g. perfectly balanced).

As a reminder, a caterpillar tree is a binary phylogenetic tree for which the induced subtree on the interior vertices forms a path graph (see Fig. 3, left). A cherry is a pair of adjacent tips on a tree. There is a single unlabeled topology for a caterpillar tree with *n* leaves. To identify the leaves conveniently, we label them using the natural ordering induced by the caterpillar tree topology. The tips in the two cherries have labels 1, 2, *n −* 1 and *n*, and the other tips are labeled accordingly (Fig. 3, left). In what follows, we use *T* to denote the caterpillar tree labeled in that manner. This labeling/ordering is not unique, but the results are independent of the labeling options for the cherries. Since we study the distribution of the transfer index *ϕ*(*χ_p_, T*) where *χ_p_* is uniformly chosen from 𝓧*_p_*, all bicolorations are equally probable, and our results remain identical with other labeling options. We call the tree *T* endowed with such labeling and coloration a bicolored oriented caterpillar tree.

**Figure 3:**
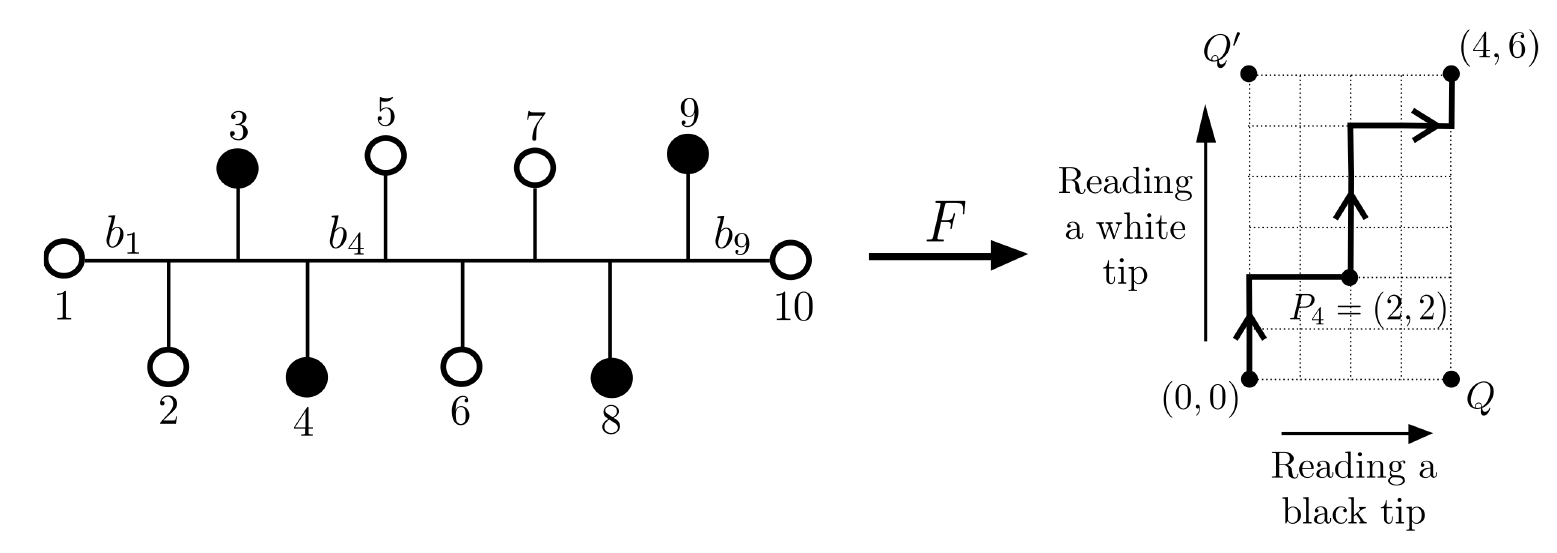
Left: a bicolored oriented caterpillar tree on *n* = 10 tips with *p* = 4; tips are numbered from 1 to 10 and branches on the path from tip 1 to tip 10 are denoted *b*_1_ to *b*_9_. Right: the associated NE lattice path from (0, 0) to (4, 6); the point *P*_4_ = (2, 2) on the path corresponds to branch *b*_4_ on the tree. Center: the function *F* that associates the tree on the left with the path on the right.

### 5.1 Correspondence between bicolored caterpillar trees and NE lattice paths

An *NE lattice path* is a path in ℤ^2^ where the only steps allowed are (0, 1) (a step towards the east) and (1, 0) (a step towards the north). From now on, we call them lattice paths for short. Let *𝓟*(*p, n − p*) denote the *p ×* (*n − p*) NE lattice, that is the set of all lattice paths from the origin (0, 0) to the destination (*p, n − p*). A path in *𝓟*(*p, n − p*) can be encoded in a single vector of length *n* indicating the sequence of steps of the path, which is an element on {*N, E*}^*n*^ with a total number of east steps equal to *p* and a total number of north steps equal to *n − p*. On the other hand, the set 𝓧_*p*_of bicolorations with *p* black tips is a subset of {**B**, **W**}^*n*^. We define the function *F* : 𝓧*_p_ → 𝓟*(*p, n − p*) that associates a lattice path to a bicolored tree by scanning the tips on the tree from 1 to *n* as follows: whenever we read a white leaf, we move towards the north; and whenever we read a black leaf, we move towards the east. Consequently, a bicolored oriented caterpillar tree on *n* tips (with *p* black tips) corresponds to a unique path in *𝓟*(*p, n − p*) and vice versa, as represented in Fig. 3. This result can be summarized as follows.

#### Lemma 10.

*The function F is a bijection from 𝓧_p_ to 𝓟*(*p, n − p*).

Let us denote the lower right corner in *𝓟* (*p, n − p*) by *Q* = (*p,* 0) and the upper left corner by *Q′* = (0*, n – p*). Observe that the two extreme paths going through *Q* and *Q′* correspond to the only bicolored oriented caterpillar trees *T* with transfer index *ϕ*(*χ_p_, T*) = 0: all black leaves cluster on one side and all white leaves on the other side. These two extreme paths can be identified with the reference bipartition *β* (Fig. 4a, left, in green). Moreover, we are able to retrieve the transfer index for any bicolored oriented caterpillar tree from the associated lattice path, as we demonstrate in the following proposition. Use *M* (*A, B*) to denote the Manhattan distance between any two lattice points *A, B* ∈ ℤ^2^, and by *M* (*γ, B*) = min_*A*_∈*_γ_ M* (*A, B*) the Manhattan distance between any lattice path *γ* and a lattice point *B*.

**Figure 4:**
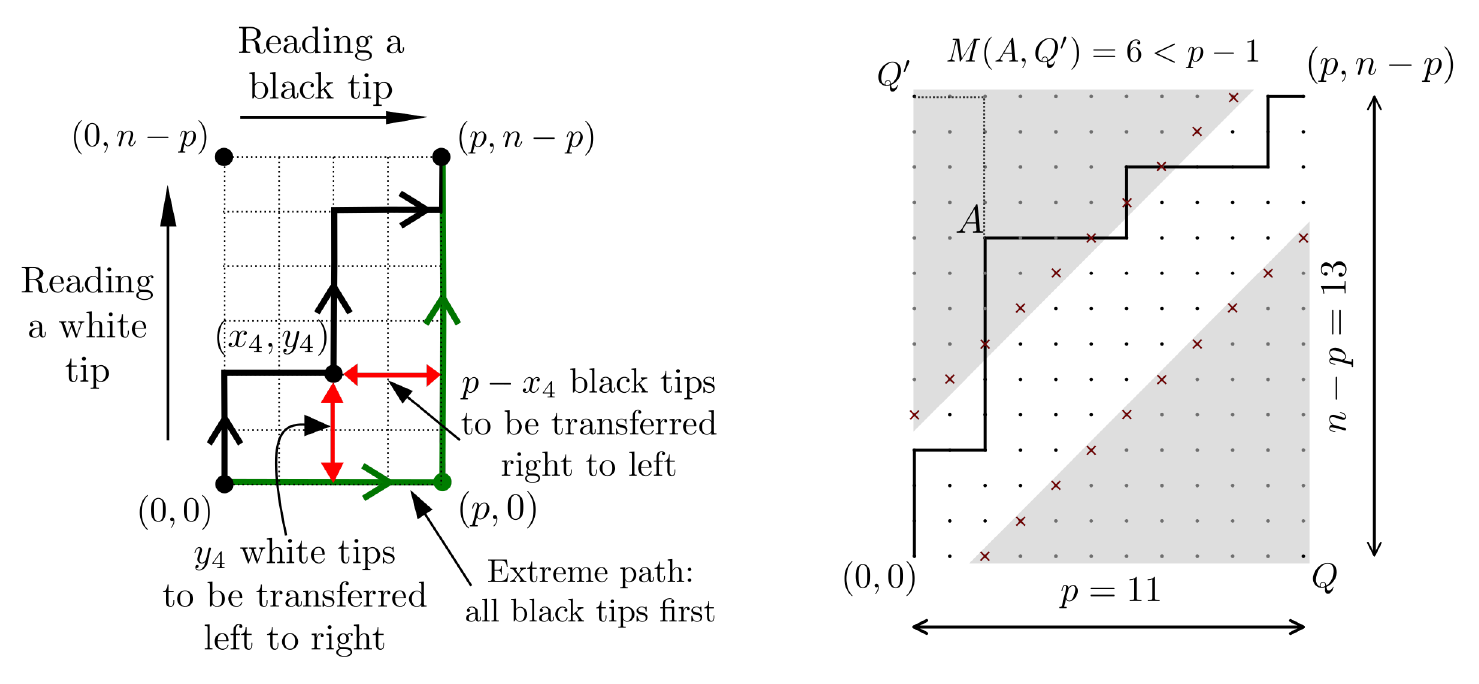
(a) Green: the extreme lattice path corresponding to the {**B**} first then {**W**} bicoloration. The branch *b* corresponding to the point (*p,* 0) in the path, results in *δ*(*χ_p_, b*) = 0. On the lattice path in bold lines, the point (*x*_4_*, y*_4_) corresponds to the branch *b*_4_ of the corresponding tree *T* with *M* ((*x*_4_*, y*_4_), (*p,* 0)) = *p −* 2 + 2 = 4 (i.e. the transfer distance of *b*_4_ as shown in Fig. 3). (b) Paths avoiding the shaded areas have a Manhattan distance to the corners *Q* and *Q′* larger than *p −* 1, and thus the corresponding bicolorations yield a transfer index = *p −* 1. When a path enters a shaded area, its Manhattan distance to the corners is less than *p* 1, e.g. point *A* is at distance 6 of *Q′* and thus the corresponding path and bicoloration have a transfer index = 6.

#### Proposition 11.

*Consider an oriented caterpillar tree T, a bicoloration χ_p_* ∈ *𝓟_p_ of its tips, and the corresponding path γ* ∈ *𝓟*(*p, n − p*)*. We have that*

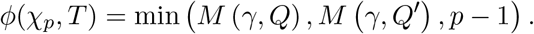

*Proof.* Consider an oriented caterpillar tree *T*, a bicoloration *χ_p_*, and the corresponding lattice path *γ* from (0, 0) to (*p, n − p*). Let us denote the *n* − 1 consecutive internal lattice points in *γ* by *P*_1_ = (*x*_1_*, y*_1_)*, …, P_n−_*_1_ = (*x_n−_*_1_, *y_n−_*_1_). Also, use *b_i_* to denote the internal branch in *T* between tips *i* and *i* + 1, for 2 *≤ i ≤ n −* 2. Lastly, let *b*_1_ and *b_n−_*_1_ be the pendant branches of tips 1 and *n* respectively (Fig. 3, left).

In the same manner that *F* associates tips in the bicolored tree with steps in *γ*, this function can be extended naturally so it establishes a one-to-one mapping from a set of branches in *T* to the interior lattice points in *γ*. More precisely, if we extend *F* to the set of branches in *T*, it holds that *F* (*b_i_*) = *P_i_*, for all 1 *≤ i ≤ n −* 1. The transfer distance between *χ_p_* and an internal branch *b_i_* is the number of tips to be transferred from one side of *b_i_* to the other side that results in the bipartition *χ_p_*. Fix 1 *≤ i ≤ n −* 1 and consider the left side of *b_i_*. Use *W* (*b_i_*) and *B*(*b_i_*) to denote respectively the number of black and white tips in this left side. By construction, we have that *W* (*b_i_*) = *y_i_* and *B*(*b_i_*) = *x_i_*, which, together with (2), leads to the following identity

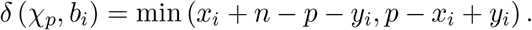

On the other hand, if we look at the Manhattan distance between the corresponding lattice point *P_i_* and the corners *Q* and *Q′*, we have that *M* (*P_i_, Q*) = *p − x_i_* + *y_i_* and *M* (*P_i_, Q′*) = *x_i_* + *n − p − y_i_* (see Fig. 4a). The same argument applies to any branch *b_i_* for 1 *≤ i ≤ n −* 1. Hence, the minimum of these distances taken over all lattice points *P_i_* in the path *γ* corresponds to the minimum of the transfer distance obtained by any internal branch in *T*, or by the leaves 1 and *n*.

Finally, all branches on the caterpillar tree that are not on the path from leaf 1 to leaf *n* are pendant branches. The minimum over all the pendant branches is equal to *p −* 1, obtained on any black leaf, so the transfer index is at least *p −* 1, as stated in the proposition. Also notice that, in the case of a bicolored cherry, the choice of the labels (1, 2 or *n −* 1*, n*) has no influence on the result since the distance from any of these pendant branches to the reference bipartition is at least *p −* 1. Since we have covered the distance obtained on any branch on the tree, we achieve the desired result.

### 5.2 Counting bicolorations through lattice paths: the transfer index distribution

Lattice paths under certain restrictions appear in various problems in probability and statistics, such as the classical ballot problem (for instance, see [10]), which leads to counting lattice paths that do not touch the diagonal *y* = *x*. Here, we are interested in a slightly different problem, but closely related to the ballot problem in the sense that we count NE lattice paths that are not allowed to touch certain boundaries.

More precisely, for fixed *n* and *p*, consider 2 *≤ l ≤ p* +1 and let *ℒ* (*n, p, l*) denote the subset of paths in *𝓟*(*p, n − p*) that do not touch *y* = *x − l* or *y* = *x* + (*n −* 2*p* + *l*). Set *L*(*n, p, l*) := *|ℒ* (*n, p, l*)*|*. The following result is from Mohanty [20]. Here, we will give a sketch of the proof that is slightly different from the one in [20], since it will be useful for understanding the upcoming results. We use ⌊·⌋ and ⌈·⌉ to denote respectively floor and ceiling functions, and 𝟙(·) for the indicator function.

#### Lemma 12

([20]). *Let c* = *n −* 2*p* + 2*l, then*

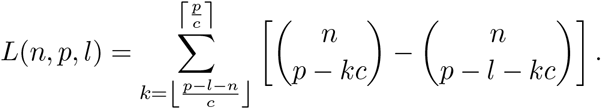

*Proof.* The proof is based on the well-known André’s reflection method [1]. First, notice that a path is uniquely defined by the *p* east steps it makes (or equivalently the *n − p* north steps), which entails that

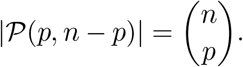

Let us count now the paths in *𝓟*(*p, n − p*) that touch the line *y* = *x − l*. If a path touching the line *y* = *x − l* is reflected from the moment it first touches this line, by switching north steps into east steps and vice versa, we obtain a lattice path ending at (*n − p* + *l, p − l*). In fact, this reflection yields a and the set *𝓟*(*n − p* + *l, p − l*), both having (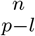) elements.

Likewise, paths in *𝓟*(*p, n – p)* that touch the line *y* = *x* + (*n – 2p + l)* can be transformed bijectively into paths in the set *𝓟*(*p − l, n − p* + *l*). Then, by applying the *inclusion-exclusion principle* [13], we have the following identity,

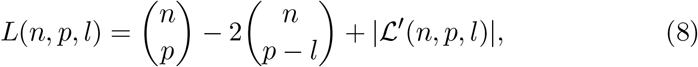

where *𝓛′*(*n, p, l*) is the set of paths that touch both lines *y* = *x − l* and *y* = *x*+(*n−*2*p*+*l*). We can then apply the reflection principle repeatedly and the usual inclusion-exclusion principle to account for paths touching both lines multiple times, leading to the above formula after some simplifications. See [20] for further details.

We can now establish the main theorem in this section.

#### Theorem 13.

*Let T be an oriented caterpillar tree and χ_p_ a bicoloration uniformly chosen from the set* 𝓧*_p_. We have for any* 2 *≤ l ≤ p* + 1 *that*

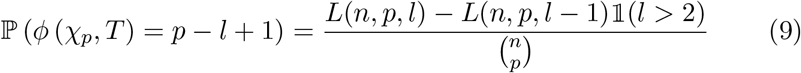

*and*

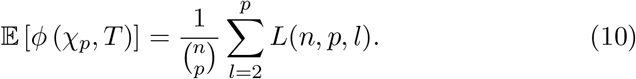

*Proof.* It is quite straightforward from the definition of *ℒ* (*n, p, l*) and Prop. 11 that for any 2 *≤ l ≤ p* + 1, the paths avoiding lines *y* = *x − l* and *y* = *x* + (*n −* 2*p* + *l*) are exactly those that remain at a distance from the corners *Q* and *Q′* greater or equal to *p − l* + 1, as it is shown in Fig. 4b, for *l* = 2. On the other hand, the paths that do touch these lines give a distance strictly smaller than *p − l* + 1. Hence, for 2 *≤ l ≤ p* + 1, a path in the set *ℒ* (*n, p, l*) *\ ℒ* (*n, p, l −* 1) (with the convention *ℒ* (*n, p,* 1) = ∅) gives a distance exactly equal to *p − l* + 1. Since the total number of paths in *𝓟*(*p, n* – *p*) is 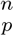 the identity (9) holds. Then, the expectation of the transfer index can be expressed as follows

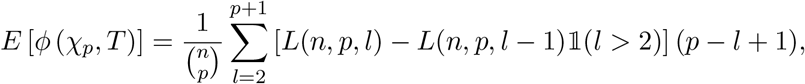

which can easily be simplified to obtain (10).

## 6 Discussion

The results we obtained in Sections 3 and 4 allow us to characterize the asymptotic behavior of the transfer support when *n* tends towards *∞*, for various growth rates of *p*, up to *p* = ⌊*n/*2⌋ and for any tree topology. However, the bounds we obtained in Section 4 are not sufficiently tight to justifiy what we observe from our simulations in Fig. 1. If we think of applications, these bounds might not be sufficient to give good estimates for the p-values of the TS distribution. We propose two conjectures that allow us to use the exact results obtained for the caterpillar tree as a proxy for the statistical significance of TS on the null models.

The first conjecture concerns the extreme case *p* = ⌊*n/*2⌋. Based on simulation results (Fig. 1 and [18]) and the proofs in section 4, we believe that for any tree topology, the expected value of TS attains its maximum over *p* at ⌊*n/*2⌋. The second conjecture (based on Fig. 1 and not shown experiments) refers to the stochastic dominance of TS for the caterpillar tree over TS for any other tree topology at *p* = ⌊*n/*2⌋.

Assuming that these conjectures (or similar ones) hold, we can bound the p-values for any *p* and any topology by the caterpillar case at ⌊*n/*2⌋, for which we have an explicit formula. For instance, consider a tree on 100 tips and a symetric reference bipartition, that is *p* = 50. The probability of this branch being supported at 50% on a caterpillar tree under the null model (the absence of phylogenetic signal) is ~ 10^*−*6^. For a tree on 20 tips and *p* = 10, the same probability is ~ 5%. These values support the idea discussed in [18] that standard levels of branch support using TBE (say *>* 70%, following Hillis and Bull [16]) cannot be observed by chance, and reveal a strong phylogenetic signal in the data, even with small trees.

For trees that are not caterpillars, deriving the distribution of the transfer index under random bicolorations appears to be challenging. It would be relevant for both theoretical and applicative reasons to characterize this distribution for a random model such as Yule or PDA, which are the most commonly used in phylogenetics [14].

## 7 Appendix: asymptotics of the transfer index on the caterpillar tree

We obtained an additional result describing the asymptotic behavior of the transfer index when *n* gets large, in the case of caterpillar trees. Similarly to our results in Sections 3 and 4, the following proposition implies that TS tends to 0 when *n → ∞* for random bicolorations. However, the speed of convergence implied by the proposition below improves the one obtained in previous sections. In particular, for *p* = ⌊*n/*2⌋Theorem 8 implies that the expectation of TS grows at most as 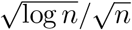, whereas for the caterpillar tree it is 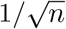 as we demonstrate below.

### Proposition 14.

*Let T be an oriented caterpillar tree and χ_p_ a bicoloration uniformly chosen from the set 𝓟_p_, for* 2 *≤ p ≤* ⌊*n/*2⌋*. The expected transfer support goes to* 0 *uniformly on p, moreover*

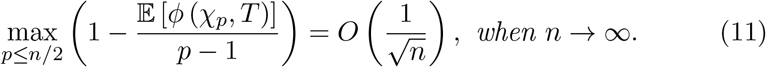

*Proof.* For any *n, p, l* with *p ≤ n/*2 and 2 *≤ l ≤ p* + 1 we have from (8) that

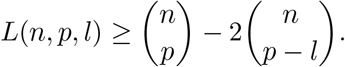

This inequality, together with (10), leads to

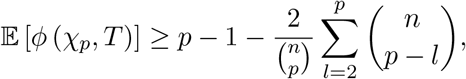

or equivalently,

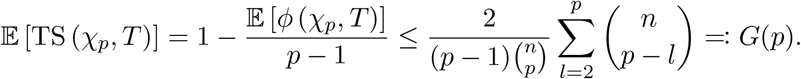

Let us now prove that *G*(*p*) is an increasing function for 2 *≤ p ≤* ⌊*n/*2⌋,

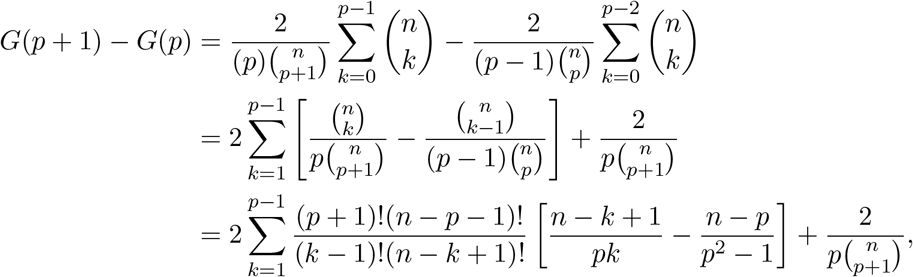

where the terms between brackets in the sum can be expanded as follows

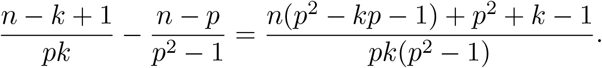

It is not hard to see that *p*^2^ *− kp −* 1 *≥* 0 for all values of *k* and *p* such that 1 *≤ k ≤ p −* 1, which implies that the previous fraction is always positive. We can then conclude that *G*(*p* + 1) *− G*(*p*) *≥* 0, so the maximum value of *G* is obtained when *p* = ⌊*n/*2⌋.

For the sake of simplicity, we suppose from now that *n* is even, but an equivalent result is obtained for odd *n* without difficulty. For *p* = *n/*2, we can use the identity 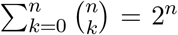 and the symmetry of the binomial coefficients 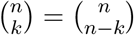 for all 0 *≤ k ≤ n,* to simplify the sum in the function *G* as follows,

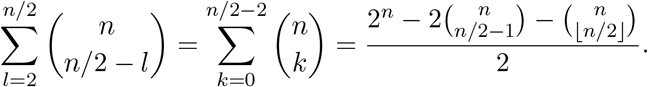

So, for all 2 *≤ p ≤ n/*2, we have that

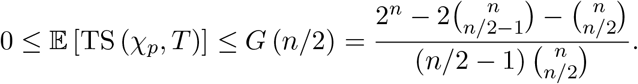

Let us now look at the asymptotic behavior of this upper bound. The well-known Stirling formula yields the following approximation for central binomial coefficients 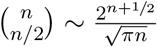 Hence

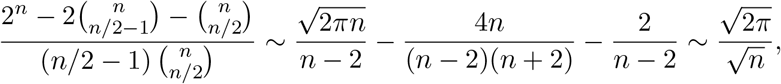

which tends towards 0 when *n → ∞*, allowing conclusion of the proof.

## Acknowledgements

This work was supported by the EU-H2020 Virogenesis project (grant number 634650 – JT, OG), by the INCEPTION project (PIA/ANR-16-CONV-0005 – MDF, FL, OG).

